# The impact of rainfall on drinking water quality in Antananarivo, Madagascar

**DOI:** 10.1101/666826

**Authors:** Alexandra Bastaraud, Emeline Perthame, Jean-Marius Rakotondramanga, Jackson Mahazosaotra, Noro Ravaonindrina, Ronan Jambou

## Abstract

Low-income cities that are subject to high population pressure and vulnerable to climate events often have a low capacity to continuously deliver safe drinking water. Here we report the findings of a 32-year investigation of the temporal dynamics of indicators of drinking water quality in the city of Antananarivo, where we assess the long-term evolution of supplied water quality and characterize the interactions between climate conditions and the full-scale supply system. A total of 25,467 water samples were collected every week at different points in the supplied drinking water system. Samples were analyzed for total coliforms (TC), *Escherichia coli* (EC), intestinal Enterococci (IE), and spores of Clostridia (SSRC). Nine-hundred-eighty-one samples that were identified as positive for one or more indicators were unevenly distributed across the series. The breakpoint method identified four periods when the time series displayed changes in the level and profile of contamination (i) and the monthly pattern of contamination (ii), with more direct effects of rainfall on the quality of supplied drinking water. The modeling showed significantly different lags among indicators of bacteria occurrence after cumulative rainfall, which range from 4 to 8 weeks. Among the effects of low-income urbanization, a rapid demographic transition and urban watershed degradation are progressively affecting the quality of supplied water and resulting in the more direct effects of rainfall events.

## Introduction

A poor capacity to provide safe drinking water, regardless of weather conditions, is of growing concern in low-income areas vulnerable to climate change [1,2]. Indeed, under global forecasts, some parts of the globe will experience increased frequency and intensity of rainfall [3] with increasing difficulties to limit storm impacts, such as flooding or increased run-off [4,5]. These events are associated with elevated turbidity [6–8] and dissolved organic matter in water sources [9], which can overwhelm treatment plans [10]. Indeed, extreme rainfall regimes are likely to be associated with drinking water contaminations [6,10], and this is predicted to be worsened by climate change [11]. Contaminated water is the main factor of diarrhea in children, and it is evident that an integrated approach to improving water supply will have an impact on the health of the population [12].

However, the relationship between rainfall patterns and water microbial quality is complex, involving an interplay between the type of water supply, the type of water source, and the treatment technology applied to water [13]. Also, susceptibility to climate change is reinforced by rapid and unplanned urbanization, poor sanitation, erosion, and low level of maintenance of the supply network [14]. Thus, the nature and the depth of the link between rainfall and water quality is not expected to be stationary. Rather, these should vary with the infrastructure and environmental changes, the time scale of study (yearly, season or day to day), and the rainfall patterns. Rainfall is a seasonal phenomenon with significant inter-annual variability [15] related to climate variations [16], extreme climatic events [6], intra-annual variability or distribution of water [17], change in duration of spells of continuous rain or no-rain events, and the total amount of water delivered during each wet spell [15]. Combined or selective impacts of these factors also depend on the catchment’s characteristics.

Due to its diverse landscape, Madagascar is exposed to a variety of weather and climate phenomena, already submitted to significant variability [18]. Antananarivo, the capital of Madagascar (Commune Urbaine d’Antananarivo - CUA), is witnessing rapid urbanization due to the arrival of 100,000 additional inhabitants per year. This evolution increases the technical constraints on infrastructure, and local services are already deficient [19]. Water supply on Madagascar only dates back since the colonial period (1952) and is outpaced by urban expansion and population growth. Consequently, home connections are still limited, and a network of about 900 standpipes supply un-piped households. In this context, storm events, heavy rainfall, and runoff can increased water turbidity and microbiological contamination [20]. Un- or insufficiently filtered, inadequately, and even adequately disinfected drinking water is thus susceptible to microbiological contamination [21]. Water utilities are required to monitor microbial indicators to assess the effectiveness of the treatment process (i.e., spores of *Clostridia*), the safety of end drinking water (i.e., *Escherichia coli, intestinal enterococci*), and the biological stability of microorganism communities in piped water (i.e., total coliforms) [21]. All changes impacting the water source, or the treatment process can affect these indicators and the drinking water quality. Characterizing the interplay between rainfall pattern and drinking water characteristics will highlight needs in monitoring and priorities in improving water production.

Due to its geographic setting, its demographic burden, and its environmental transition, Antananarivo is a suitable example for setting up new strategies to survey water treatment based on predictive mathematic models. Predictive models could also be used to adapt the activities of dispensaries based on the burden of diarrhea. First, we analyzed how the water quality had evolved over the last 30 years, and then how the rainfall pattern has impacted long-term water quality in the supply system under environmental and technical shifts. We also focused on the current week-to-week relationship between rainfall patterns characteristics and water quality to determine the main factors regulating water contamination.

## Materials and Methods

### CUA and its drinking-water supply system

#### Site of study

Antananarivo is located in the central highlands of Madagascar at 1,300 m altitude (18°55’ S latitude and 47°32’ Longitude). The city is nestled among twelve hills and lies in the natural floodplain of the Ikopa River, which skirts the city to the south and west (Fig. 1). The river and its tributaries play an important role in rice-dominated agricultural production. The metropolitan area spreads over 220 km^2^, with an estimated population of 3,058 million inhabitants in 2018 [22]. This area currently experiences significant challenges due to flooding during the rainy season.

**Fig 1.**
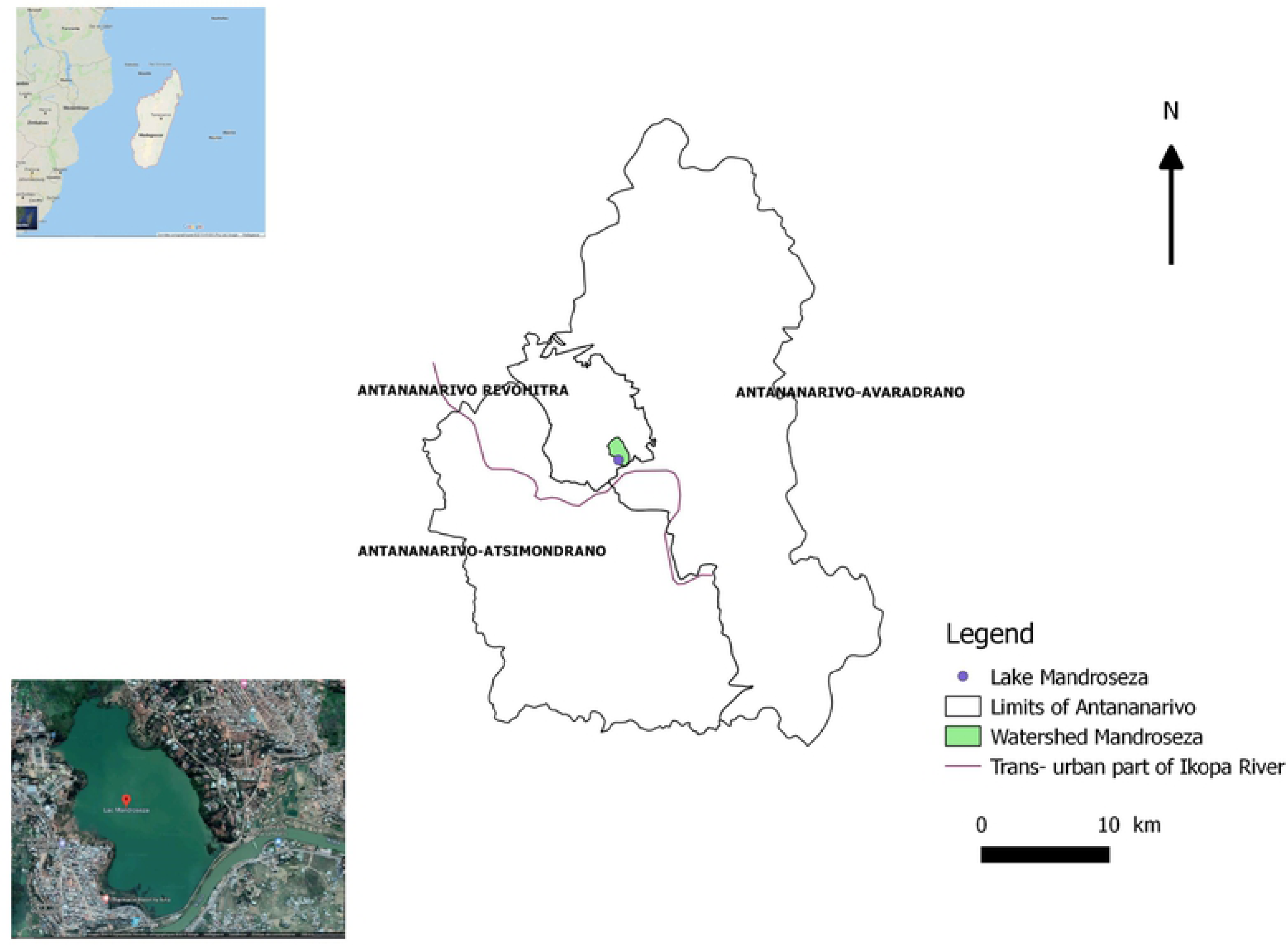
Location of Mandroseza Lake inside the limits of Antananarivo.

#### Climate

Antananarivo experiments a subtropical highland climate, warm and temperate classified as Cwb by the Köppen-Geiger system [23]. Summers are rainy, with most of the precipitations falling in December, January, and February. The winters are dry, especially in June, July, and August. The dry season occurs from May to September (i.e., from week no.14 to no.40), and the wet season from November to April (i.e., from week no.41 to no.13). From 1985 to 2017, the annual average of rainfall was above 1500 mm with extremes in January (above 300 mm) and June (less than 10 mm). The city of Antananarivo has experienced cyclones over the past 20 years, including Geralda in January 1994, Giovanna in February 2012, and Enawo in January 2017. These induced severe flooding as in February 2015 and 2016.

#### Water supply

In 2015, according to the National Water and Electricity Utility (JIRAMA), the whole drinking water supply system represents 1000 km of pipes for 80,000 subscribers (supply rate 56.8%). Water is provided by the Ikopa River, whose flow is diverted to the 1.41 km^3^ artificial Mandroseza Lake from where water is pumped [24]. Two water stations, Mandroseza I and II, with a daily production of 93,000 and 62,000 m^3^ per day (m^3^/d) respectively, supply 30 reservoirs. The treatment process includes coagulation, flocculation, decantation, filtration, chlorination, and neutralization steps [25].

### Data collection

#### Water sampling and analysis

Four different points of the network (e.g., piped households, administrative buildings, standpipes) are randomly investigated daily, 5 days per week. Each sample is collected in 500 ml sterile containers with 10 mg sodium thiosulfate and stored at 4 to 10°C until processing at the Institute Pasteur laboratory within 18 to 24 h. Microbial water quality is assessed by counting negative and positive samples for several microbial indicators, including *Escherichia coli* (EC) and Total Coliform count (TC) [26,27]; intestinal enterococci (IE),[28,29]; spores of Sulfite-Reducing Clostridia (SSRC) [30]. Between 2010 and 2016, the 100 ml filtration method was preferred for testing SSRC [31]. Since 2014, the IDEXX Quanti-Tray methods have been implemented for testing IE [32], EC and TC [33]. Criteria for a negative sample is set to “undetectable microorganism in any 100 milliliters (ml)” [34] and in any “20 ml for SSRC”, before 2010 and after 2016 [30].

Temporal patterns of microbial water quality are expressed as the frequency of positive samples reported to the total number of samples collected during the period (monthly and weekly) over 32 years (from 1985 to 2017, except 2009 due to the insurrectional crisis). Monthly series had 6.41% missing values, and weekly series had 18.69% missing values. For multivariate analysis, missing data were replaced by the median of the related month for monthly series and of the related week for weekly series.

#### Rainfall data

From 2007 to 2017, monthly rainfall was collected from the data from *Direction G*é*n*é*rale de la M*é*t*é*orologie*. Daily rainfall was collected from the IRI-International Research Institute for climate and society. To have the same time-step for rainfalls and contamination, these data were summarized as cumulative rainfall by month or by week.

### Statistical analysis

#### Breakpoint detection method

To detect specific periods or obvious trends within contamination markers and rainfall time series, we applied a breakpoint detection method implemented in the Strucchange R package [35]. We used the method of simultaneous estimation of multiple breakpoints proposed by Bai and Perron in 2003 [36]. The method was run using the default parameters of breakpoints function, and the number of periods was estimated by minimizing the BIC (Bayesian Information Criterion).

#### Fourier analysis

To test if the variable “month” affects the contamination markers and rainfall pattern, we applied a Fourier transform to each variable using the TSA (Time Series Analysis) R package [37]. The computed periodograms from this transformation were tested. If there is a month effect in the time series, the periodogram should have a peak at time 12 (corresponding to 12 months). The significance of the amplitude of the periodogram at time 12 is tested using a permutation test (i.e., comparison with a random sequence with a significance level at 0.05) [38]. All p-values are available in the supporting information (S1 Table).

#### Hierarchical clustering

To check for similar current profiles of contamination, we focused on data from the last period provided by statistical analysis of contamination change. We applied a hierarchical clustering algorithm with Euclidean distance and Ward distance. The clustering was applied to the four contamination markers (IE, EC, TC, and SSRC). Total contamination was not used for clustering to avoid collinearity with IE, TC, EC, and SSRC. Rainfall and total contamination were added to the graphical representation for interpretation.

#### Auto-regressive integrated moving average (ARIMA) models

To investigate the specific relationship between drinking water contamination and rainfall, we have run three models following: a “naive” model (i) that consists of forecasting the contamination of a given week by the mean of the previous corresponding weeks. This model does not account for the effect of rainfall and is used as a benchmark for further comparisons: any prediction model achieving higher prediction error is not relevant. Two different ARIMA models are fitted on each marker: an ARIMA model, with no extra covariate (ii); and an ARIMA model adjusted on cumulative rainfalls of the previous weeks (iii), with a shift varying from 1 to 10 weeks. The optimal number of cumulative weeks is estimated by minimizing the prediction error (root mean square error, RMSE) assessed by cross-validation in the years 2016 and 2017. A likelihood-ratio test was used to compare the goodness of fit of statistical models. All parameters of ARIMA models are automatically selected using a stepwise procedure which minimizes the BIC, implemented in the auto.arima function of forecast package [39]. The Portmanteau test was used to conclude that no residual autocorrelation remained in the models [40]. This procedure allows investigation of the effect of rainfall and cumulative rainfalls on contamination markers; it also allows for the quantification of the number of cumulative weeks optimal to predict contamination.

## Results

### Contamination of the water over the 32 years

A total of 971 samples have been identified as positive for one or several microbial indicators among 25,467 samples collected over 32 years (365 months). This accounts for 3.8% of non-compliant samples, unequally distributed over time. Indeed, during the period from 1989 to 2004 (175 months = 75% of the months), no contamination was reported.

### Breakpoints in the yearly pattern of contamination

The time series of monthly water contamination frequencies showed significant shifts over the years for all indicators (IE, EC, TC, and SSRC). Change points and the associated 95% confidence intervals for total contamination (IE, EC, TC, and SSRC), as well as rainfalls, are summarized in Table 1 (Breakpoint detection method).

**Table 1.**
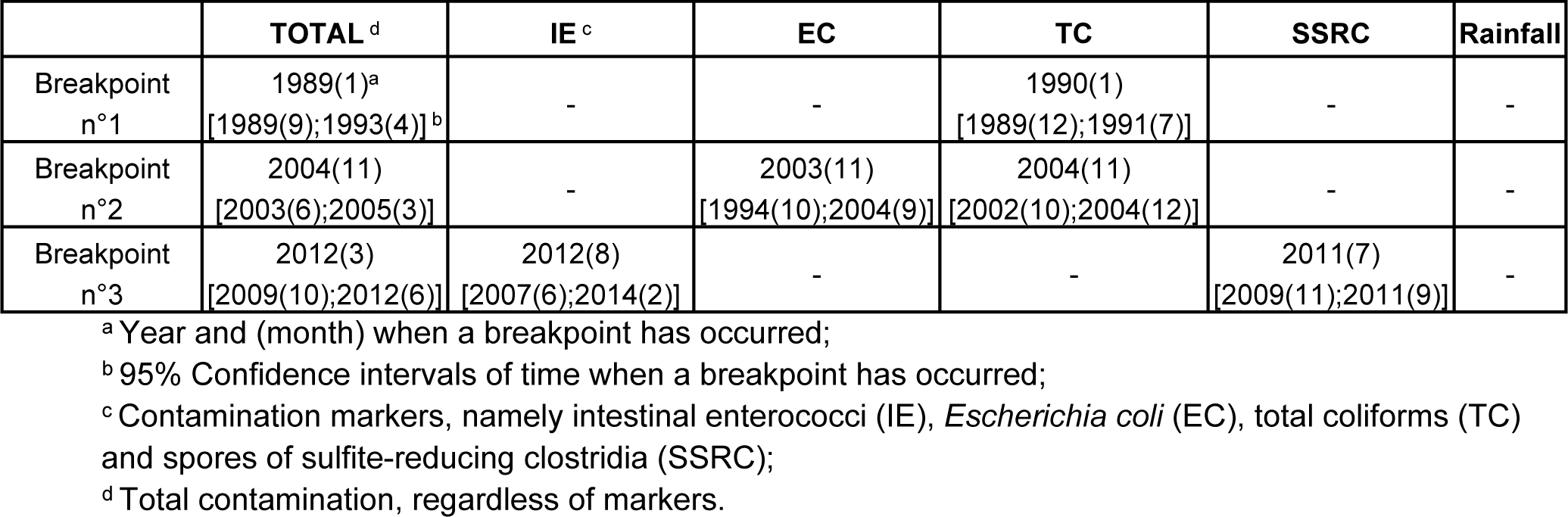
Breakpoints in contamination markers and rainfall time series.

Three breakpoints occurred over the “total contamination” series, defining four periods: i) before 1990, ii) between 1990 and 2005, iii) between 2005 and 2012, and iv) after 2012. Time series analysis also captured specific change-points for each specific contamination marker [i.e., 1990 and 2004 (TC); 2003 (EC); 2011 (SSRC) and 2012 (IE)]. For these periods, the monthly average of contaminated samples varied from 1.1% (the intermediate period from 1990 to 2005) to 9.5% (the last 5 years) (Fig 2). Periods 1 (from 1985 to 1990) and 3 (from 2005 to 2012) showed similar levels of contamination (4.7% and 4.2%). While the period from 1990 to 2005 was relatively free-from water contamination. The period from 2012 to 2017 showed a significant decrease in drinking water quality. Over the 30 years, no breakpoint or obvious trend was detected in the rainfall.

**Fig. 2.**
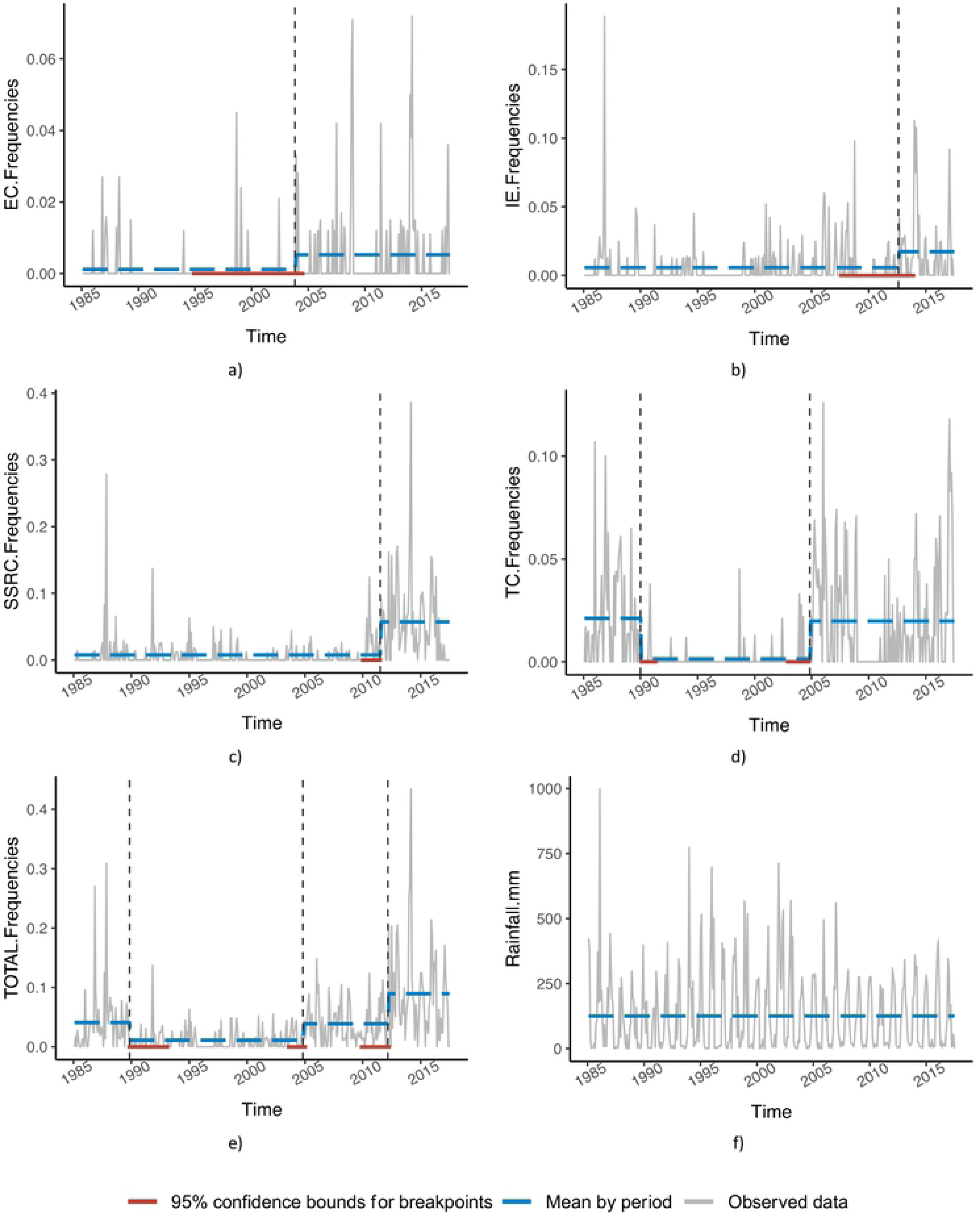
Time series of drinking water contamination frequencies in Antananarivo’s (Madagascar) water supply and rainfall from 1985 to 2017, using the period from breakpoints method. The time series are displayed in grey. The periods are represented by dashed vertical black lines; the mean of the time series within each period is indicated by a dashed blue line. Confidence intervals associated with changepoints are shown as red lines.

Over the whole series, and except for the poorly contaminated 1990–2005 period, SSRC contamination events have continuously increased (Fig 3d), with the recent median of contamination events reaching 4.8% (Fig 3k). During the last period, IE contamination also increased, with median contamination events rising from 0 to 0.8% (Fig 3k). EC contaminated samples remained sporadic throughout the periods, with medians of the periods close to zero. However, the means of contamination increased very slightly from 0.1% (Fig 3e) to 0.7% (Fig 3k) (breakpoint in 2004). For TC, three out of four periods showed baseline contamination events with half of the months harboring 1% of contamination.

**Fig 3.**
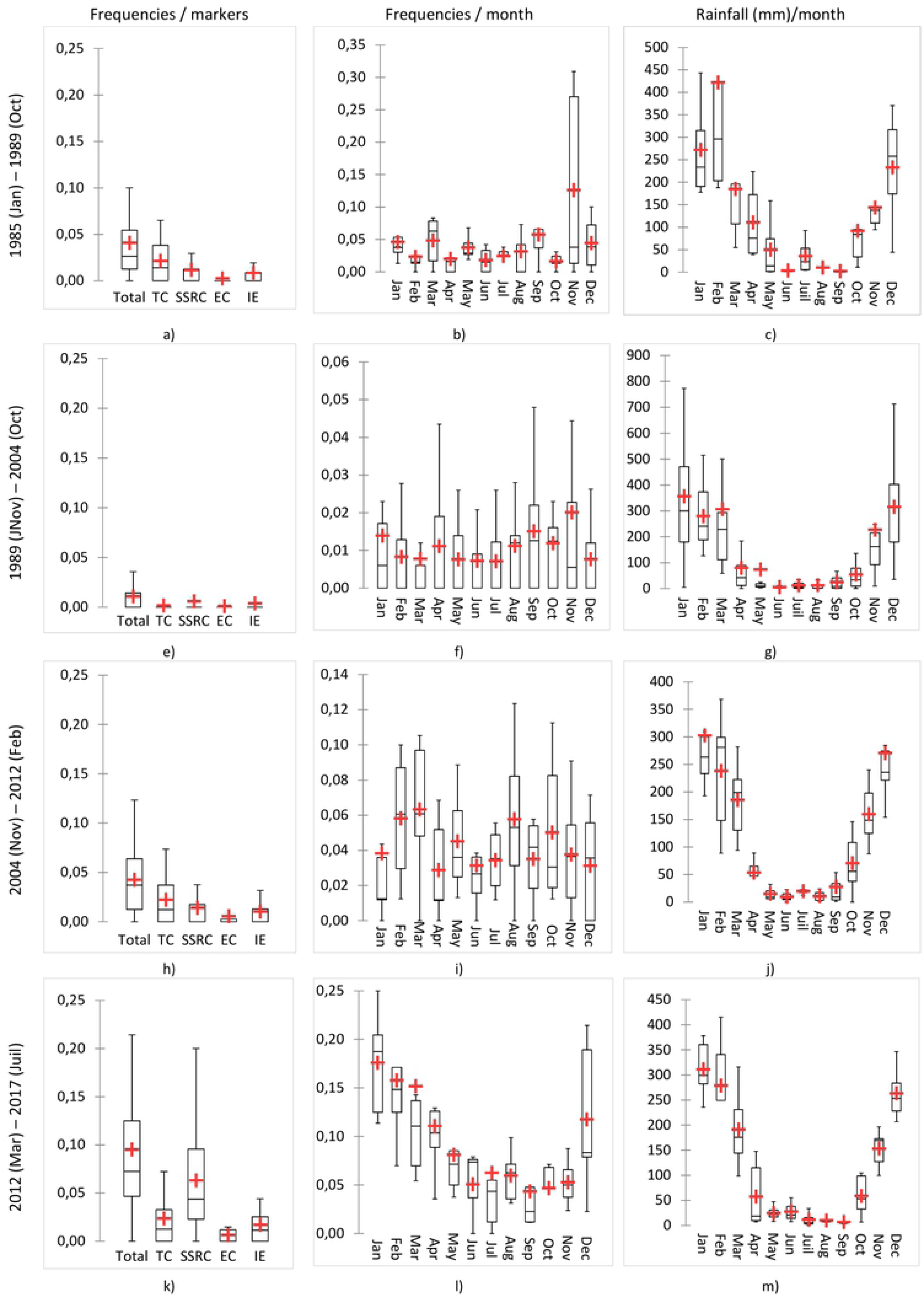
Distribution of drinking water contamination frequencies in Antananarivo’s (Madagascar) water supply and rainfall from 1985 to 2017. Box plots are displayed with a mean (red cross).

### Change in monthly pattern of contamination

As the months harboring the highest contamination events also varied over time (Fig. 3), the month effect had to be tested independently from the periods. The periodogram method and permutation test showed significant 12 months periodicity in data of several years for TC, IE, and rainfall (p-value < 0.05) (S1 Table). This means that TC and IE contamination events and rainfalls have preferentially occurred at a specific month during the year, whatever this month. For SSRC, the test was barely significant (p-value ∼ 0.05), suggesting that it suffers from a lack of power to detect a specific pattern for this parameter. No month effect was detected for EC (p-value ∼ 0.3 > 0.05).

More than 13 years ago, November harbored the maximum of sample contamination accounting for the highest mean of the period, with 12.6% (Fig 3b) and 2% (Fig 3f), in the first and second period, respectively. In Antananarivo, November is also the month of the first heavy rainfall (Figs. 3c, g, j, and m). During the following periods, contamination events progressively slipped preferentially at the beginning of the year, with means and medians of contaminated samples reaching 6% and 17% respectively in March (third period) and January (last period) (Fig. 3i and l). This is in accordance with the rainiest months (Fig. 3j and m). During the last 6 years, contamination events spread over the December to April period (Fig. 3l). Thus, monthly contamination distribution can be superposed to rainfall distribution (Fig. 3l and m).

## Relationship between water contamination and rainfall

### Profile contamination clustering

Correlation between the percentage of contaminated samples collected for one month and the rainfall measured during the same month was first searched using a clustering strategy conducted on the whole data set collected from March 2012 to the current date. For each month, rainfall, total contamination, IE, EC, TC, and SSRC were considered as variables for multiple component analysis (MCA) and automatic ascendant classification. The months grouped in a cluster and exhibited a similar profile of contamination. Figure 4a displays the clustering tree. Four clusters were determined according to the level of each contamination marker (IE, EC, TC, and SSRC). The scatter plot displays the distribution of all markers within each cluster. Rainfall and total contamination were also displayed (Fig 4b). Cluster 1, the largest, included 34 observations characterizing to months with low contamination in the context of low rainfall. Cluster 4, the smallest, included seven observations that demonstrated high contamination in TC and IE/EC, in the context of middle rainfall level. Clusters 4 and 3 exhibit similar total contamination. However, cluster 3 reported higher SSRC, lower TC, and IE/EC contamination and higher rainfall than cluster 4. This suggests that a similar total contamination rate might be associated with different contamination profiles. Cluster 2 demonstrated high contamination for all markers in the context of high rainfall.

**Fig 4.**
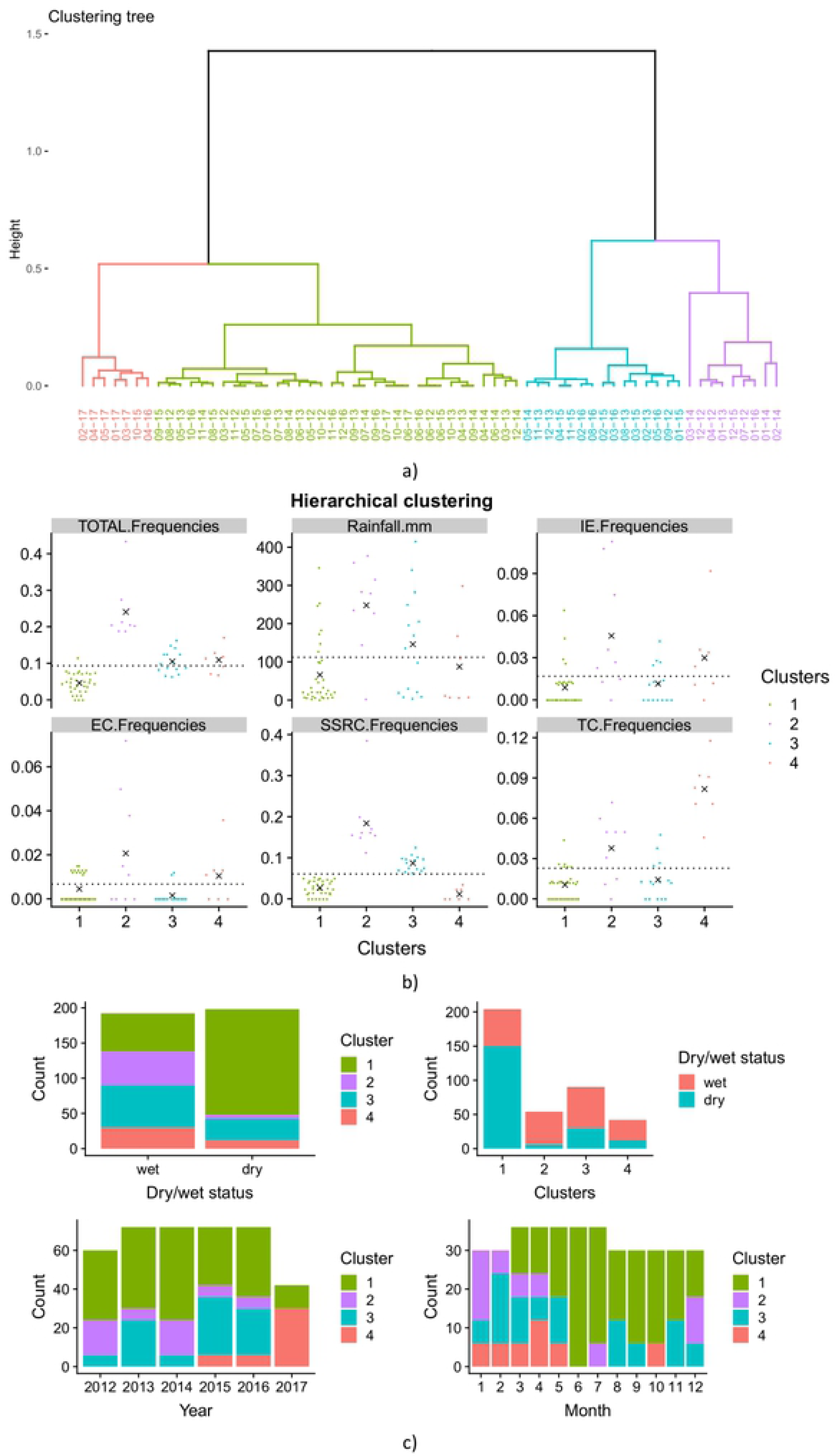
Hierarchical clustering of monthly observations from drinking water monitoring in Antananarivo’s (Madagascar) water supply from 2012 to 2017. a) Clustering tree; b) Scatter plot displays the distribution of all markers within each cluster. The black crosses are the mean within the cluster of the corresponding variable. The dotted black line is the overall mean. c) Bar plots explore the repartition of each cluster by wet/dry seasons, by year and by months.

In summary, during the dry seasons, the level of contamination was relatively low and mainly caused by SSRC (cluster 3). During the wet seasons, contamination was related to all other contamination markers (cluster 2) or by SSRC (cluster 3). Contamination by TC and IE/EC (cluster 4) was also more reported during the wet season.

The contamination profile also varied over the years. TC and EI/EC contaminations emerged three years ago and became predominant in 2017 (cluster no.4). Whereas in 2013, 2015, and 2016, contamination was mainly sustained by SSRC (cluster no. 3) and in 2012 and 2014 by all other markers (cluster no. 2). At the month level, contamination profiles varied cyclically according to the seasons. The wettest months of January and February showed high contamination, mainly caused by all types of microorganism (cluster no. 2) and SSRCs (cluster no. 3) respectively. This high contamination level occurred until May, regardless of the marker. Except during these first 5 months of the years, low or no contamination was observed, suggesting a rainfalls effect on emergence and persistence of contamination.

### Rainfall and contamination modeling

According to previous data, the cumulative effect of the amount of rain fallen over previous weeks on contamination can be suspected. However, this effect could vary with the markers. For each marker, three models (observed means, ARIMA model with no covariate, and ARIMA model adjusted to the optimal number of cumulative precipitations) were compared to quantify the impact of cumulative rain. The values fitted by the three models were reported (Fig 5), and determinants of best models were summarized in Table 2.

**Table 2.**
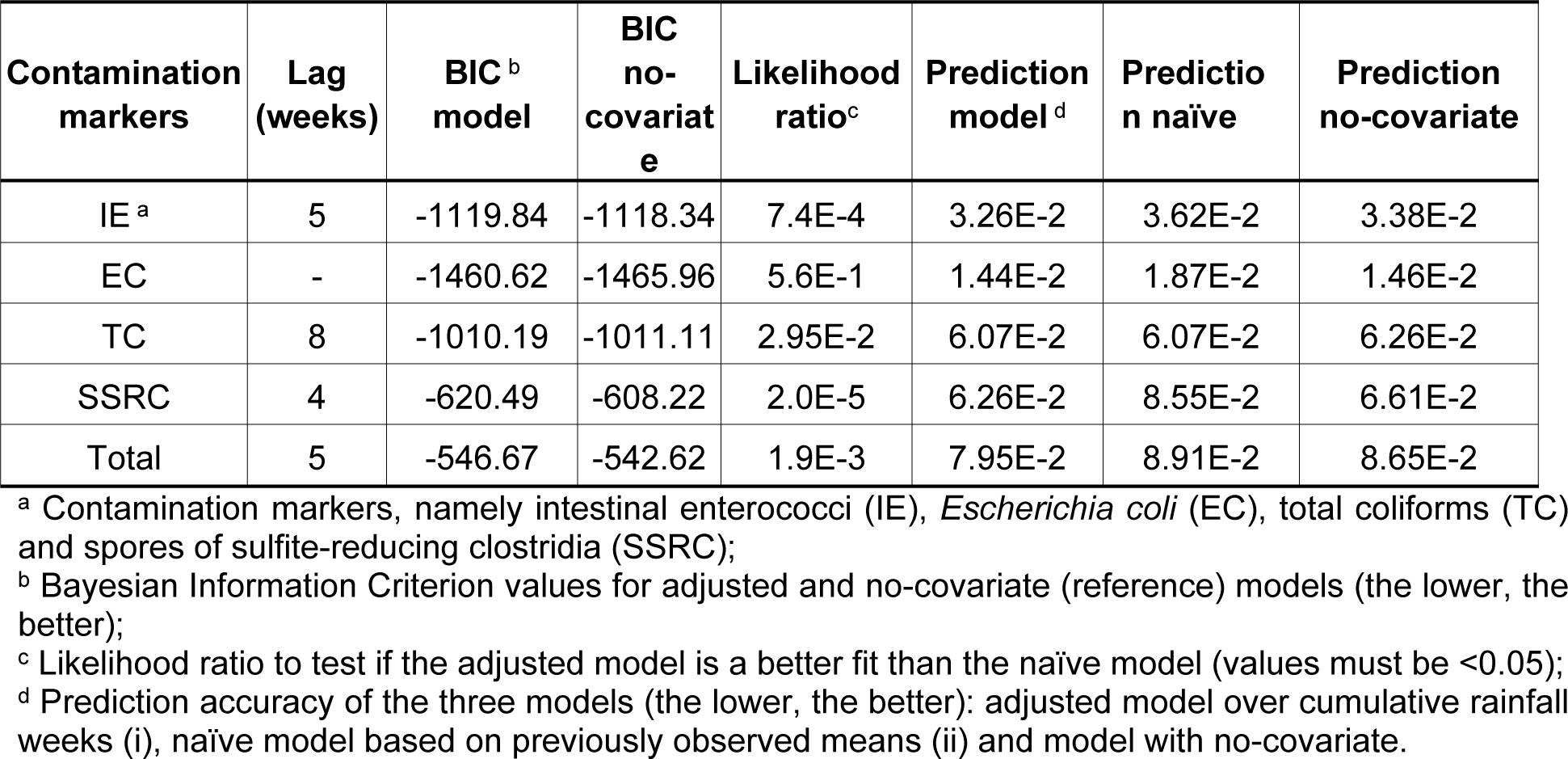
Determinants of best ARIMA model adjusted to the optimal number of cumulative rainfalls.

**Fig 5.**
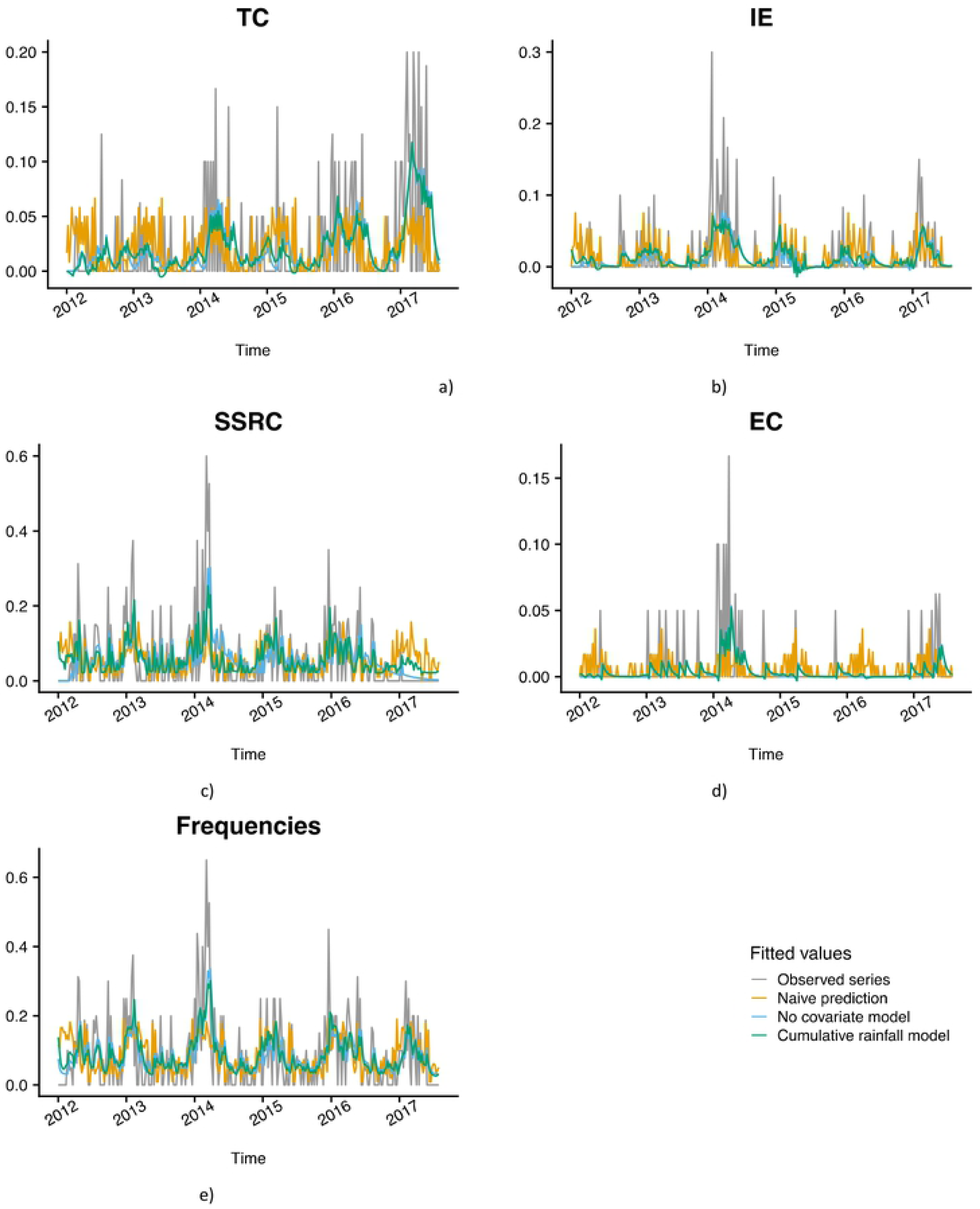
Modeling drinking water contamination in Antananarivo’s (Madagascar) water supply from 2012 to 2017. The following figure shows the observed series in grey, the values fitted by the mean in orange, by the ARIMA model with no covariate in blue, and by the ARIMA model adjusted on the optimal number of cumulative rainfalls in green.

The ARIMA model adjusted with cumulative weekly rainfall was found as the most accurate with the lowest BIC value and with a significant ratio test of likelihood (p-value <0.05). The Portmanteau test concludes that no residual autocorrelation remained in the models (p > 0.05 for all models).

In summary, apart from EC, these models showed that weekly cumulative rainfalls had an impact on drinking water quality with different time lags according to the contamination markers. For total contamination, a lag of 5 weeks of cumulative rainfall led to the best model (BIC = −546.67) when comparing to the model with no covariate (BIC = −542.69, a p-value of the likelihood ratio test = 1.9e-03<0.05). Prediction performance is also better than the other models (7.95E-2 < 8.91E-2 and 8.65E-2).

For other specific markers and in agreement with MCA, the different lags suggest the chronological emergence of contamination markers after the weekly rainy periods: 1) SSRC contamination events generally occurred first after 4 weeks of cumulative rainfalls; 2) IE contamination events occurred after 5 weeks of cumulative rainfalls; 3) TC emerged last, within 8 weeks of cumulative rainfalls. For EC, this procedure estimated that 3 weeks was the optimal cumulative rainfall (BIC = −1460.62). Nevertheless, the likelihood of the model does not significantly increase compared to the model with no covariate (BIC = - 1465.96, a p-value of likelihood ratio test = 5.6e-01 > 0.05). This suggests that the procedure is not able to detect how rainfall affects EC rate. Prediction error of the model adjusted on 3 weeks of cumulative rainfall (∼1.44e-02) does not lead to an improvement of the prediction accuracy of the naive model (∼1.87e-02) nor of the ARIMA model with no covariate (∼1.46e-02). This could be due to a lack of power of the model as EC contamination events were too sporadic (9% of the series).

In conclusion, the emergence of SSRC, IE, and TC are differentially linked to cumulative weekly precipitations, but no significant impact of cumulative rainfall could be detected for EC.

## Discussion

### Drinking water quality issues

Drinking water supply systems are commonly affected by the raw water quality and the capacity of the treatment process to remove pollutants. The biological and physicochemical stability of the water in the network is also a quality issue, as secondary contamination of the water can occur in the supply networks [41]. Among these factors, the effectiveness of the treatment process is certainly the most important and must be carefully adapted to the quality of the incoming raw water. However, hydraulic conditions (e.g., flow velocity and intermittent water supply) and water quality (e.g., presence of sediments or excess of turbidity) can enhance bacterial growth and sediment dissolution in final drinking water [17,42].

The drinking water quality guidelines issued by the World Health Organization serve as an international reference point for national or regional regulations and standards on water safety. These are not mandatory but must be adapted to the technical and environmental specificities of each country [43]. Global contamination markers include at least *Escherichia coli* (EC), an indicator of recent fecal contamination. Out of the 43 microbiological parameters identified as markers of contamination, Madagascar has implemented monitoring of three parameters in addition to EC, namely total coliforms - TC (i), intestinal enterococci – IE (ii), and clostridium spores - SSRC (iii). TC are mainly bacteria capable of surviving and growing in water. Directly measured after treatment, TC is an indicator of treatment effectiveness. Otherwise, TC can also be used as an indicator of network hygiene. Intestinal enterococci are indicators of fecal pollution that survive for a long time in water and multiply in certain ecological niches, such as sediments or aquatic flora. Monitoring of Clostridia is used as an indicator of the filtration process. Clostridia can persist for a long time in pipes, attached on sediments [44]. Clostridia are present in oxygen-free niches, such as soil habitats, sewage, rivers, lakes, and the intestinal tract [45,46]. In Madagascar, only spores of Clostridia are monitored.

The contamination events can be punctual or persistent over time, depending on the strength and durability of the impact of the factor(s) on water supply. Environmental and water resource permanent changes can lead either to major failures in the drinking water production system or only to a lasting depreciation of the quality of the water supply. These changes occur at different times, so long time series are unlikely to be homogeneous. Indeed, study of the quality of the water over the last 30 years in the city of Antananarivo highlights four distinct and unequal periods. A common point in these four periods is the low proportion of samples contaminated with EC, which indicates that, regardless of the ecological context or the technical performance of the treatment plan, the station still can remove recent fecal contamination [47].

### Water supply and demand issues: Signal of imbalance

According to 30 years of data, the 1985–1993 and 2004–2012 periods seem to illustrate the baseline level of water quality supplied by the station. Studies must now focus on periods during which improved or deteriorated water quality was registered to analyze factors leading to these anomalies. Since the quality of the water supplied has significantly improved over a 15-year period, with contamination events now rare, we searched for changes that could have occurred in the treatment plan, and that would improve water quality. Changes in the design of the drinking water production system of the Antananarivo city can first explain most of this improvement.

Indeed, in Antananarivo, the water production system dates to 1921 and the Mandroseza I plant, still operating, dates to 1957. To date, the plant was producing 100,000 m^3^ per day [25]. These infrastructures are under severe pressure from rapid and unplanned urbanization; more than 120,000 new urban dwellers arrive per year, and there has been a global population increasing from 252,000 inhabitants in 1960 to 2 million in 2010. The average population growth over the last 30 years has been 4.9% [22], and the current yield of water supply (volume sold / volume produced) barely reaches 60% [48]. Facing this increase of the urban population, a second plant was built (Mandroseza unit II in 1993), which coincided with a long phase (1989–2004) of reduction in contamination events (particularly from total coliforms). Exceeding production capacity and aging of filtration systems seem to be the most important factors having affected Mandroseza’s water treatment. Indeed, filtration is an import factor in preventing coliform proliferation, and its improvement can reduce coliforms by a factor of three [49]. This 15-years period constituted an interlude between two periods (1985–1989 and 2004–2012).

We detected a resurgence in TC contamination after 2004. This might be related to the beginning of a new period of increase in the gap between the nominal and actual production capacity of water treatment. With a 160,000 m^3^ production per day, the plant was progressively overloaded, and the balance was, once again, broken [24,50]. In March 2017, a new subunit (Mandoseza II bis) was created to increase the capacity from 3,000 m^3^/h to 3,900 m^3^/h, but it is too soon to assess the impact of this on water quality. However, Antananarivo is expected to host nearly 3,400,000 inhabitants in 2020. Urbanization and demographics will again affect the equilibrium between water supply and demand. The critical point will the economic capacity to again set-up new infrastructures and to find a new economic model for water supply [51].

### Ecological disruptions and damage to the water resource

Despite climate changes, there are no clear trends in rainfall data. The climate of Antananarivo is nevertheless marked by a two-headed distribution of rainfall (end and beginning of the year). Drought periods are much more pronounced in the highlands and extend from April to October. Precipitation during the first months of the year is expected to increase [52], which would increase the vulnerability of the water supply system during this period. Also, Antananarivo has experienced several sequential extreme weather events since 2012, experiencing high contamination events throughout the period. The occurrence of Cyclone Giovanna at the end of February 2012 coincides with the beginning of the period with the highest contamination rates. The rains induced by Cyclone Felling in January 2013 contributed to the rise in water levels. The February 2015 rainfalls triggered floods and surging flood waters breached several dams around the capital, Antananarivo. The heavy showers had been plaguing the island since the end of 2014 were followed by the passage of Cyclone Chedza in January 2015. Since December 2015 – and more specifically in January 2016 – Antananarivo has also experienced torrential rains leading to significant damage. Enawo, which was a category four cyclone, struck Antananarivo in March 2017.

Correlation analyses for cumulative rainfalls and water contamination confirm that rainy periods are more susceptible to contamination and that the occurrence of the different markers is chronological. In the 4 to 8 weeks following heavy rainfall, the contamination markers typically emerge in the order: 1) SSRC, 2) IE, and 3) TC. Contamination of the water network is, therefore, a cumulative phenomenon or a chain reaction that starts at the beginning of the rainy season. The emergence of SSRC without other contamination markers might indicate soil leaching [53] during the first rains in November, which will gradually overload the station after 4 weeks.

The basin is degraded and vulnerable to erosion phenomena [54,55], with sediment flow impacting the efficiency of filtration. Also, the soils of the highlands are poorly protected by vegetation (deforestation). Subjected to heavy rains, these are sites of very strong erosion [56,57]. The Ikopa River watershed is, therefore, severely impacted by erosion. In the highlands, those cities that use surface water also have events of drinking water contamination, mainly driven by the SSRC [58]. Failures in the treatment system appear to have occurred, especially during wet periods.

The first IE emerge at week 5 after the beginning of heavy rainfalls. These IE indicate environmental contamination, persist for a long time in the system and are resistant to chlorine. Their presence is usually associated with a loss of efficiency of the filtration systems. IE colonize ecological niches such as sediments, lakes, or plants, and contamination could result from a deterioration in the quality of the resource or its environmental context [59,60]. This increase in IE was only seen during the more recent period. Indeed, in 1995, the silting of the Ikopa River was estimated at 81 m^3^ per year per km^2^ of the watershed, and sediments concentrated mainly upstream of dams (particularly Mandroseza dam) [9,61–63]. Since then, the depth of Lake Mandroseza had gradually decreased from 7.5 m to 3 m and has begun to be invaded by non-aquatic plants. Demographic and ecological changes have also occurred, including deforestation of watersheds and disturbance of the protection perimeter related to urbanization. The Mandroseza basin has increased from 30 to 50 hectares of the urbanized area [64]. Since then, although EC contamination events are not significant (median zero), the range of contaminated samples has increased. Nevertheless, the emergence of EC contamination is not significantly related to cumulative rainy weeks. The treatment system can limit the occurrence of EC, even though urban runoff can be heavily loaded with this bacterium [65].

After 8 weeks of cumulative rainfall (January–February), TC appeared. These contamination events are delayed and not directly related to precipitation. Unlike SSRC and IE, these contamination events seem related to another parameter. This event could be the accumulation of sediments or the deterioration of the cleanliness of the supply networks. The loss of filtration efficiency also creates conditions for the proliferation of TC in the supply network [21,66].

### Bias induced by changes or lakes in monitoring

Over the last 30 years, some parameters have been ignored during water quality monitoring. Sampling turbidity measurements, for example, have only been processed and recorded by the laboratory since 2016. Similarly, the evaluation of organic matter in the water network (the simplest being the determination of permanganate oxidation) would also be necessary to assess hygiene. Monitoring of the chlorine level should allow the characterization of the response to this phenomenon. The chlorine demand resulting from the difference between the amount of chlorine added and the residual chlorine in the system tends to increase as the hygiene conditions of the system deteriorate. It is also likely that the disruption in 2012, which was characterized by a high level in SSRC, was related to the change in method and volume (× 5) for the measurement of SSRC (NF T 90- 415 vs. NF EN 26461-2). Apart from this case, no testing changes impacted on fecal contamination (EC and IE) and TC, which were highly stable during this period.

## Conclusions

- The bacteriological quality of the supplied water in Antananarivo has gradually deteriorated in recent years. The rapid demographic transition has gradually weakened the treatment plant by maintaining an imbalance in production capacity and water requirements. Treatment efficiency is currently affected by overproduction, and exponential unplanned urbanization or low yield water production were aggravating factors.
- The degraded environmental context of watershed and water resource recently affect the raw water quality and reinforce the impact of heavy rainfall on drinking water quality. Siltation of lake resources and erosion were aggravating factors during rainy periods, introducing contamination markers attached to sediments into the supply system (i.e., spores of sulfite-reducing of Clostridia and intestinal enterococci).
- The overload of the filtration system mainly occurs between 4 and 5 weeks of cumulative rainfall favoring strong contamination in January and February. Regrowth conditions of bacteria are evident with the emergence of total coliforms after 8 weeks of cumulative rainfall. Consequently, the vulnerability to persistent contamination and biological instability generally persist during rainy periods. Nevertheless, the treatment process is sufficient at removing *Escherichia coli*.
- The implementation of the treatment plant in 1993 had a long and positive impact on drinking water quality, mainly in decreasing contamination events by total coliforms. An appropriate filtration treatment upgrade strategy could be effective in managing the microbiological quality of the water in the supply system. Otherwise, a fair balance between the duration of filtration (flow rate) and the quantity of available treated water must be found.
- Stability in testing methods and expansion of monitoring parameters are needed to better assess changes of the interplay between climate and environmental or technical context of water supply.

## Acknowledgments

We are grateful to Dr. Philippe Cecchi for methodological guidance and The Center of Bioinformatics, Biostatistics, and Integrative Biology (C3BI) of Institut Pasteur at Paris for research assistance.

## Supporting information

**S1 Table. P-values from permutation test for testing yearly periodicity in contamination time-series data**.

